# Of puzzles and pavements: a quantitative exploration of leaf epidermal cell shape

**DOI:** 10.1101/361717

**Authors:** Róza V. Vőfély, Joseph Gallagher, Grace D. Pisano, Madelaine Bartlett, Siobhan A. Braybrook

**Author notes:** These authors contributed equally to this manuscript (Róza V. Vőfély, Joseph Gallagher).

## Abstract

**Summary:** The epidermal cells of leaves lend themselves readily to observation and display many shapes and types: tabular pavement cells, complex trichomes, and stomatal complexes^1^. Pavement cells from *Zea mays* (maize) and *Arabidopsis thaliana* (arabidopsis) both have highly undulate anticlinal walls and are held as representative of monocots and eudicots, respectively. In these two model species, we have a nuanced understanding of the molecular mechanisms that generate undulating pavement cell shape^2–9^. This model-system dominance has led to two common assumptions: first, that particular plant lineages are characterized by particular pavement cell shapes; and second, that undulatory pavement cell shapes are common enough to be model shapes. To test these assumptions, we quantified pavement cell shape in the leaves of 278 vascular plant taxa and assessed cell shape metrics across large taxonomic groups. We settled on two metrics that described cell shape diversity well in this dataset: aspect ratio (degree of cell elongation) and solidity (a proxy for margin undulation). We found that pavement cells in the monocots tended to have weakly undulating margins, pavement cells in ferns had strongly undulating margins, and pavement cells in the eudicots showed no particular degree of undulation. Indeed, we found that cells with strongly undulating margins, like those of arabidopsis and maize, were in the minority in seed plants. At the organ level, we found a trend towards cells with more undulating margins on the abaxial leaf surface vs. the adaxial surface. We also detected a correlation between cell and leaf aspect ratio: highly elongated leaves tended to have highly elongated cells (low aspect ratio), but not in the eudicots. This indicates that while plant anatomy and plant morphology can be connected, superficially similar leaves can develop through very different underlying growth dynamics (cell expansion and division patterns). This work reveals the striking diversity of pavement cell shapes across vascular plants, and lays the quantitative groundwork for testing hypotheses about pavement cell form and function.

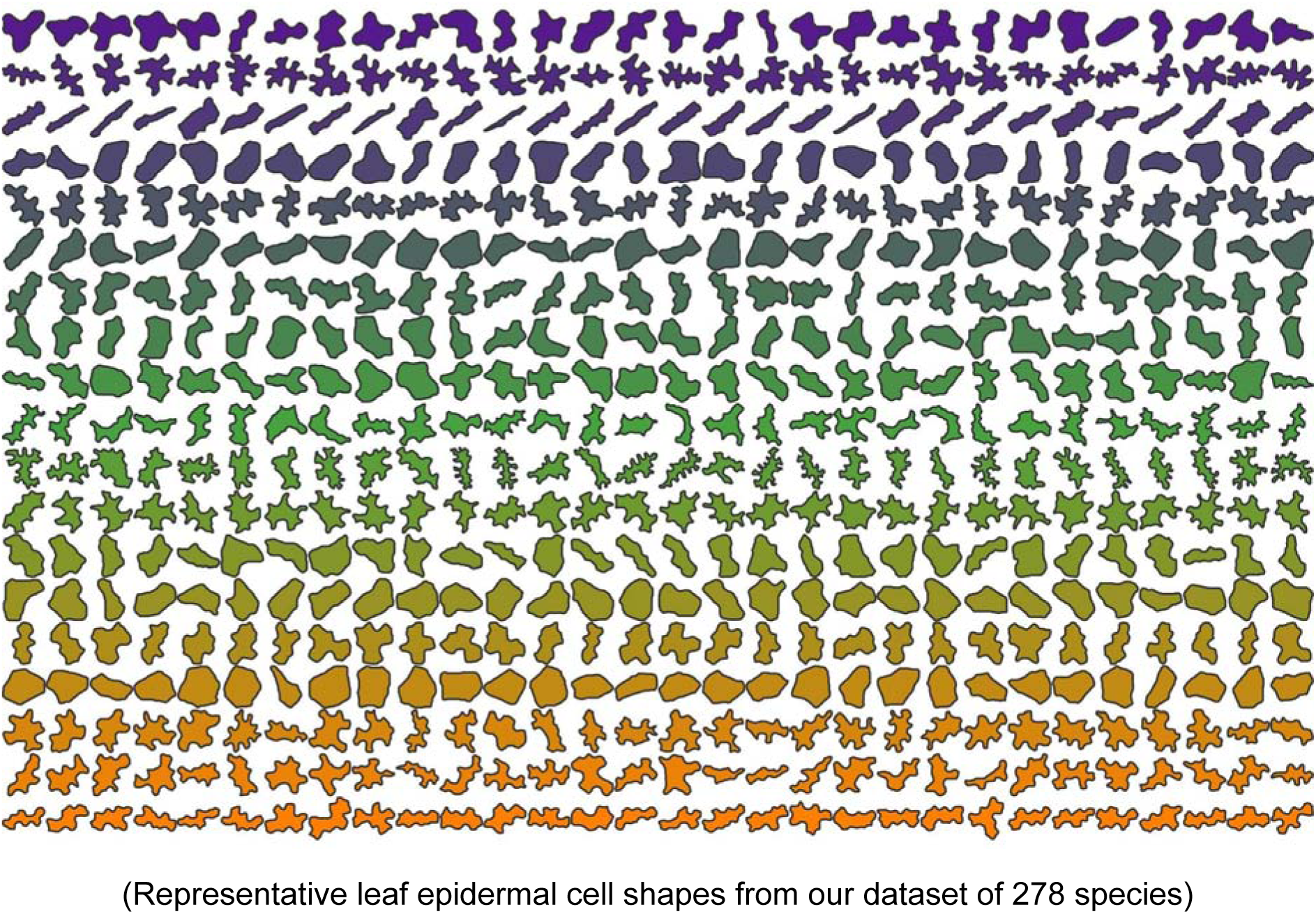

## Introduction

The first cell was described by Robert Hooke in 1665; the empty cells of sectioned cork, seen under a microscope, were likened to the cells of a honeycomb^10^. Since that time, scientists have been observing plant cells in all of their diversity of form. The epidermal cells of leaves lend themselves readily to observation and display a great diversity of shapes and types: tabular pavement cells, complex trichomes, and stomatal complexes^11^. Pavement cell shape, in particular, has been the focus of many recent studies, probing the mechanistic basis of cell shape generation ^2–6,8,12–14^.

Molecular studies of pavement cell shape generation have focused almost exclusively on model genetic species such as arabidopsis, maize, and *Oryza sativa* (rice) ^6,12,15^. All of these epidermides present dramatically undulating cell margins, while maize and rice (both grasses) exhibit extreme cell elongation. From such studies a molecular framework for pavement cell shape generation has been proposed which explains extreme undulation as a result of differential cell wall properties underlain by differential cytoskeletal patterning^13^. Although intensely studied at a molecular level, and despite an early qualitative survey of leaf pavement cell shape^16^, it remains unclear how common margin undulation is in the plant kingdom.

Another abiding mystery is the biological reason (if any) for margin undulation – how does margin undulation affect organismal form and function? There are two long-standing hypotheses in the field: (1) undulations may increase cell-cell contact between adjacent cells, allowing for more efficient chemical signalling^17^; (2) undulating margins may increase epidermal integrity (think of a zipper)^7^; and (3) undulations may help leaves flex^18^. A third, more recent hypothesis proposes that larger, isotropic, cells undulate to alleviate the stress caused by their own growth dynamics^19^. This cell-strength hypothesis^19^ was put forth on the basis of observations in species closely related to arabidopsis, and therefore represents a much-needed foray into non-model species. However, a phylogenetic context is an important consideration for any experimental designs of this type. Without taking phylogeny into account, one cannot be sure whether observed correlations are for functional reasons, or because of underlying relatedness of the species under study^20^. Quantitative assessments of cell shape, coupled to modern phylogenetic methods, allow for this disentangling of contingency and functional relevance.

Here, we present a broad quantitative survey of epidermal pavement cell shape, analysed in an explicitly phylogenetic context. Utilising morphometric methods, we determined two useful metrics for describing margin undulation (solidity) and base cell shape (aspect ratio) across a wide swathe of the plant kingdom. We mapped solidity and aspect ratio values onto a phylogenetic tree of ferns and seed plants, and tested for phylogenetic signal. Phylogenetic signal assesses the propensity for trait values to be similar between closely related species. We found that while particular cell shape metrics characterized the ferns, gymnosperms, and monocots that we sampled, we could only detect phylogenetic signal at shallow phylogenetic levels in the eudicots. Our results indicate that cell shape is extremely diverse across the land plants, particularly in the eudicots, and that the mechanisms driving the development of plant cell shape should be explored in systems beyond the current dominant model systems.

## Materials and Methods

### Sampling

Fully expanded adult leaves were collected from healthy plants grown in one of two locations between September 2015 and December 2017: The Botanic Garden of the University of Cambridge or the UMass Amherst Natural History Collection (see Table S1 for full species list). Note that cultivars and wild taxa were analysed together in this study.

### Sample preparation

Two methods of sample preparation were used; First, when possible, epidermal peels were removed from the adaxial side of the leaf. When this was not possible, the abaxial side was attempted. Secondly, when peels were unachievable a dissection and maceration protocol was followed. In detail, roughly 5×5 mm asymmetric trapezoids were cut from the leaves, near the midrib, halfway along the length. The asymmetric shape allows keeping track of adaxial and abaxial sides through the several-day-long process. These pieces were placed in multi-well plates and soaked in ~ 1 ml of a 1:7 mixture of acetic acid and 100% ethanol overnight at 4°C, stirred at 50 rpm. The following day the solution was removed and samples were washed three times for 10 minutes. After the last wash, water was replaced by 1 ml of 1M NaOH solution and left to stand for 24 h at room temperature, without stirring. Following this, the samples were washed again as before, and the solution was replaced by 1 ml of a solution containing 250 g chloral hydrate dissolved in 100 ml of a 1:2 mixture of glycerol and water. The samples remained in this solution for 3-5 days, until they became fully transparent. When the clearing finished, the samples were washed again as before and stored in water. Note that gaps in joint adaxial/abaxial sampling resulted from temporal shifts in methods as well as technical challenges of peeling in some cases.

### Staining and Imaging

Samples were stained with 0.1% toluidine blue in water overnight, mounted on glass slides and covered with a coverslip. Images were acquired at 100X, 200X, 400x, 700x or 1000x magnification (depending on what was found appropriate for a given sample) using a Keyence VHX-5000 digital microscope (Cambridge; Keyence UK & IL) or an Axioplan microscope (Amherst; Zeiss, DE). Whenever possible, images were taken from both sides of the sample, at the same magnification. Images were saved in .tif format.

### Segmentation

Automatic segmentation of these images proved to be very difficult due to image defects on different length scales: dust grains, trichomes and hairs, uneven staining, varying light intensity across the image. Some of these can be eliminated by simpler image processing methods (filtering, smoothing) but others cannot. Therefore, we chose to perform segmentation manually, using a freely available image editor (GIMP^21^) using a tablet PC and stylus, resulting in a black-and-white image of cells. The outlines of the cells were extracted in MATLAB^22^ using basic built-in functions. For each species, 30 cells were segmented per side (when both available).

### Leaf shape

Leaves were flattened and scanned in front of a white background at a resolution of 300 dpi. These images were first binarised using an automatically determined simple threshold and the outlines were then extracted using MATLAB. One leaf sample (or leaflet for compound leaves) per species was used, the same leaf from which cells were extracted.

### Shape processing and statistical analysis

Cell outlines were used to calculate traditional morphometric descriptors (absolute area in μm^2^ for cells and mm^2^ for leaves, aspect ratio, circularity and solidity) and to extract the elliptic Fourier composition. Calculations were done using the momocs^23^ package in R. Aspect ratio was calculated by calculating the ratio of and outline’s width to length (See Fig. 1; coo_width/coo_length). Circularity (coo_circularitynorm) was defined as: perimeter squared divided by area, normalised by 4*π*. Solidity was calculated by dividing the area of a shape by the area of its convex hull (see Fig. 1; coo_solidity). For the Fourier analysis of cell shapes 20 harmonics was utilized based on a cumulative harmonic sum >99.9% and test fitting outlines with undulatory margins (Fig. S3). A normalized Elliptical Fourier Analysis was performed using momocs (efourier_norm, for area); normalization was included as randomly sampled species, when stacked, exhibited clean alignments without rotational artefacts.

**Figure 1.**
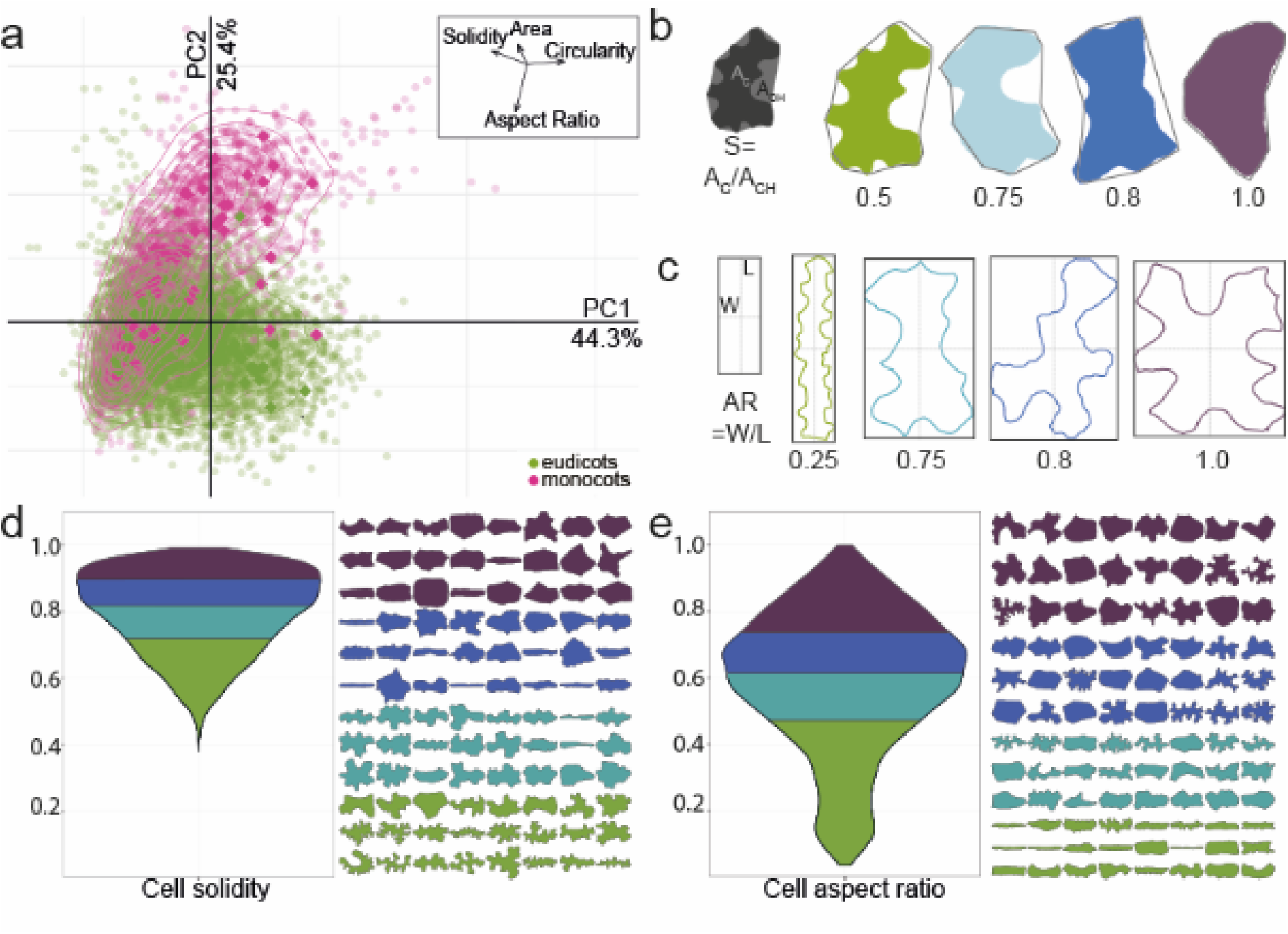
Traditional shape descriptors describe variation in base cell shape and margin undulation. (a) A Principal Component Analysis of all epidermal cells sampled from monocots (pink) and eudicots (green) using traditional shape descriptors of aspect ratio (AR), area (A), circularity (C), and solidity (S). In this analysis, 69.7% of shape variance in the data set was explained by the first two principal components. The vectors describing the morphospace (inset) demonstrate how each shape descriptor relates to the first two components. (b) An illustration of cell solidity (S) calculation as the ratio of cell area to the convex hull area and its results from four representative cells with constant aspect ratio; colouring of representative cells matches quartiles bellow in d. (c) An illustration of aspect ratio (AR) calculation as the ratio of maximal width to maximal length and its results from four representative cells with constant solidity value; colouring of representative cells matches quartiles bellow in e. (d) The distribution of solidity values for our entire data set, coloured according to quartiles. 24 cells from the median of each quartile are displayed with the same colour coding for reference. (e) The distribution of aspect ratio values for our entire data set coloured according to quartiles. 24 cells from the median of each quartile are displayed with the same colour coding for reference.

Principal component analysis was performed on the full dataset, again utilising the momocs package. Data from all cells were used in PC analysis presented here, not means or representative cells. For cells, PCA was conducted using all cells from all species. For leaf shape, the outline from one leaf (or leaflet for compound leaves) per species was used to calculate traditional morphometric descriptors as above. Correlations between traditional metrics (Spearman’s rho) were examined in R.

### Tests for phylogenetic signal

The data matrix for phylogenetic analysis was constructed by extracting sequences from the matrix used for inferring a recent megaphylogeny of vascular plants^24^. Where there was not an exact match for the species we sampled, we selected another species in the same genus from the megaphylogeny matrix. When there was no genus match, we retrieved sequences for each of the missing species from Genbank. The megaphylogeny matrix includes 7 gene regions. We aligned each of these gene regions individually using MAFFT, as implemented in Geneious, and concatenated each of these regions into a single matrix. A constraint tree, including all taxa in our analysis, was extracted from a megaphylogeny of vascular plants using phylomatic^25,26^. We used a constraint tree because we were not trying to infer phylogenetic relationships, but instead to generate a tree (with branch lengths) that we could use in downstream analyses. Model and partitioning scheme selection was performed using PartitionFinder. We analyzed our data matrix under the maximum likelihood information criterion using RAxML, as implemented on the CIPRES webserver^27,28^.

The resulting phylogeny was used in tests for phylogenetic signal using the R package PhyloSignal^29^. Phylogenetic signal is the tendency of traits in related species to resemble each other more than in species drawn at random from the same tree. In a test for phylogenetic signal, the null hypothesis is that the values of a particular trait are distributed independently from their phylogenetic distance in a tree. There are a number of tests for phylogenetic signal; we selected local Moran’s I which is designed to detect local hotspots of positive and negative trait autocorrelation^29,30^. The phylogeny figure was generated using the R package ggtree^31^, with final editing performed in Illustrator (Adobe).

### Data Availability

The datasets generated during the current study, including the phylogenetic datamatrix and trees, have been deposited to dryad (dryad link to follow). Mean shape metrics are included in Supplemental Table 1. For some species, scanning electron micrographs are also available upon request, although not included in this study. Code for analyses and figure generation is available at XXXXX (Bartlett lab’s github).

## Results and Discussion

### Most vascular plants have slightly elliptical pavement cells with weakly undulating margins

To survey pavement cell shape across vascular plants, we sampled leaf epidermides from 278 –vascular plant species, taking current phylogenetic hypotheses into account_^32,33^. To quantify cell shape, we used the traditional shape descriptors of area, circularity, aspect ratio, and solidity (Fig. 1; See Methods for definitions). We utilised these traditional metrics because we found that elliptical Fourier analysis did not perform well with our extremely diverse dataset (Fig. S1); elliptical Fourier analysis did a reasonable job of capturing aspect ratio variance but not margin undulations (Fig. S1). In a principal components analysis (PCA) with the traditional metrics, the sum of PC1 and PC2 together accounted for 69.7% of shape variance (Fig. 1a, monocots and eudicots as an example; Fig. S1, all clades). The vectors describing the traditional morphospace indicated that aspect ratio and solidity were strong perpendicular separators of cell shape (Fig. 1a, inset). Solidity was calculated by finding the area of the cell shape and dividing it by the area of the convex hull (Fig. 1b); the convex hull of an object can be conceived of as a rubber band stretched around the perimeter, so that in undulating cells the convex hull gaps away from the true perimeter (Fig. 1b). To calculate aspect ratio, cells were oriented according to their longest axis and the longest cell width was divided by the longest cell length in this orientation (Fig. 1c). Circularity represents how deviant a cell shape is from a perfect circle^7,34^ and captures both margin undulations and aspect ratios deviating from 1. This merged property was illustrated by the morphospace vector for circularity which was the inverse sum of that for aspect ratio and solidity (Fig. 1a, inset). Thus, we concluded that solidity and aspect ratio were good descriptors of margin undulation and base cell shape, respectively.

To determine whether pavement cells across vascular plants were characterized by a particular base cell shape or undulation pattern, we examined solidity and aspect ratio across our sampling. We found that most plant species displayed weak margin undulation. Solidity values for all species sampled occupied a range between 0.38 and 1, with a median of 0.802 (Fig. 1d; Table S1). This skew indicated that while most sampled pavement cells showed some degree of undulation, a minority of species sampled displayed complex margins (low solidity). Both arabidopsis and maize pavement cells fell within the bottom 8% of solidity values for seed plants (S_At_ = 0.67, S_Zm_ = 0.63). The solidity metric is imperfect: curved cells with simple margins will also have a lower solidity value due to the calculation of convex hull area (Fig. 1b). In addition, solidity describes the deviation of the perimeter from the convex hull, but it doesn’t provide information on the pattern of that deviation. For example, a margin might have a few deep lobes or many shallow lobes but have similar solidity values. This may have also been an advantage in our analysis: when the pattern of lobing was variable within a species (e.g. Arabidopsis) solidity would have been less sensitive to small variances in lobe number; note that in a single species context, a new modification of Fourier analysis would prove an excellent tool to assess such variation^35^. Our analyses of cell aspect ratio indicated that while most pavement cells were mildly elliptical in their base cell shape (median > 0.5); highly anisotropic or truly isotropic cells were rare in our data set (Fig. 1e). The distribution of aspect ratio across all species sampled occupied a range of 0.069 to 0.805 with a median value of 0.643 (Fig. 1e; Table S1). Thus, we found that the average epidermal cell in plants might best be represented by a slightly anisotropic cell with weak margin undulation.

Highly undulate pavement cells are not common (Fig. 1d) and as such our molecular models of shape generation require modulation to reflect the diversity observed in the plant kingdom. The current molecular model for undulation (or protrusion) formation in arabidopsis has actin concentration at positions of protrusion outgrowth and microtubule bundling restricting growth across indentations. This role of actin in protrusion outgrowth is consistent in maize and rice^6,15^. Patterns of actin and microtubules in several other species with undulating cell wall margins are also consistent with this model, although microtubules likely have numerous roles in pavement cells^4,12^. Given the distant relationship between arabidopsis, a core eudicot, and maize and rice, core monocots, this mechanism may be common to all monocots and eudicots. In arabidopsis, the patterning of alternating actin/microtubule patches is set up by active RHO-RELATED PROTEIN FROM PLANTS 2 (ROP2). Active ROP2 promotes RIC4-mediated fine actin accumulation while suppressing RIC1-mediated microtubule bundling^3^. In a situation where protrusion number and depth vary quantitatively on a phenotypic continuum (Fig. 1e), it is possible that the alternating pattern of actin and microtubules (and their controlling RICs) may be distinct between different species. It is equally probable that the patterning is conserved but the wall components and modifiers differ, leading to different wall mechanics and growth.

Differential cytoskeletal patterning also likely leads to differential wall thickness and material composition, as recently shown in several species with undulating pavement cell margins^18^. Differential biochemistry and mechanics of the wall are likely contributors to cell shape formation in arabidopsis^36^. These differential material properties must also be considered when considering the ‘reason’ for undulation: the mechanical integrity of tissues during stretching may be important^18^. It has recently been proposed by modelling cell stresses that undulation helps an individual cell deal with its geometrically imposed stress as it gets isotropically larger^19^. A small sampling of plant species (n=16) showed a positive correlation between cell area and cell ‘lobeyness’^19^ (‘lobeyness’ was solidity calculated as a perimeter ratio, as opposed to area ratio here). In line with this hypothesis, pavement cell margin undulation is used in reconstructing paleoclimates because larger shade leaves often have larger pavement cells with more undulate margins^37,38^. Thus, we tested for this in our dataset, but found no correlation between mean cell area and mean solidity (Fig. S2). Differential wall mechanics might explain this lack of correlation in our broad sample: there may be multiple ways to be strong. For example, making the cell walls thicker or materially more rigid in a larger cell could also compensate for geometrically-imposed stress. An analysis of wall composition, thickness, and mechanics in large cells with varying degrees of undulation would prove interesting.

### Patterns and diversity in pavement cell shape by vascular plant clade

To examine if trends in base cell shape and margin undulation might exist across the major clades of vascular plants with true leaves (megaphylls^39^), we examined the distributions of aspect ratio and solidity in the following clades: ferns, gymnosperms, the ANA grade, magnoliids, monocots, and eudicots^32,40^. We found that the ferns displayed a shift towards more undulate cells on average (Fig. 2a). Fern pavement cell margins have been described as more undulating than in the eudicots^41^, an observation that holds generally true in our data set; however, ferns exhibited the widest range of solidity values (0.38 to 0.98; Fig. 2a). The distribution of solidity values within the eudicots was also broad, although slightly less so than in ferns (Fig. 2a). Monocot and gymnosperm pavement cells tended to exhibit higher solidity values (less undulating margins) consistent with the qualitative literature for monocots^16,41–45^(Fig. 2a). With respect to aspect ratio, the ferns, early diverging angiosperms, and eudicots displayed normal distributions centering between 0.6 and 0.7, representing the slightly ellipsoidal base shape norm (Fig. 2b). In gymnosperms and monocots, the distributions were more skewed with medians below 0.4 indicating a trend toward a more anisotropic base shape in these groups (Fig. 2b). Taken together, these results indicate that pavement cells in the ferns, monocots, and gymnosperms share specific aspect ratio and solidity traits, while pavement cell shape has diversified in the eudicots.

**Figure 2.**
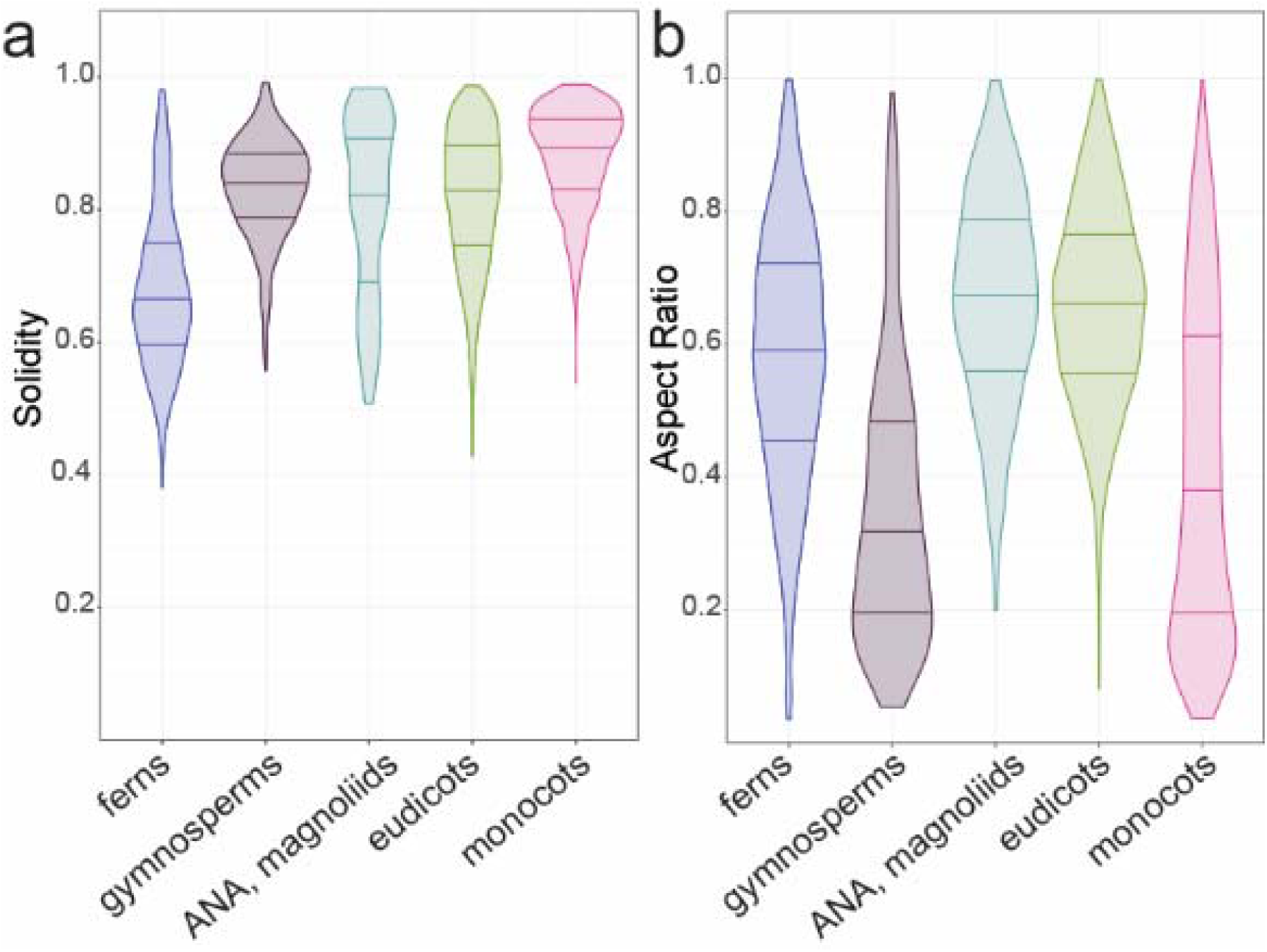
Solidity and aspect ratio distributions vary between clades. (a) Solidity and (b) aspect ratio data are presented as distributions by clade for ferns, gymnosperms, the ANA grade and magnoliids, monocots, and eudicots. Note that the mean solidity values for both arabidopsis (SAt= 0.67) and maize (SZm=0.63) fall within the tails of the eudicot and monocot distributions, respectively. No sample size scaling has been applied.

Our initial analysis did not take phylogeny into account, and cannot detect signal in specific orders or families obscured by considering, for example, ‘eudicots’ as a single group. To account for phylogenetic relationships, we mapped cell solidity and cell aspect ratio values onto a phylogeny of all the species that we sampled and tested for phylogenetic signal. Although related species tend to resemble one another, this is not true for every trait in every lineage. Tests for phylogenetic signal assess whether particular traits are more similar between closely related species than between distantly related species, or between species drawn at random from the same phylogenetic tree^30,46^. Most of the ferns we sampled (n=31/35, 89%) showed evidence for phylogenetic signal for solidity, with more complex cell margins (low solidity, Fig. 3a). In contrast, most core monocots (n=38/46, 82%) have cells with less complex margins, falling within the first two quartiles of solidity (values closer to 1; Fig. 3b). Similarly, many core monocots had a strong signal for highly anisotropic cells (low aspect ratio, n=23/35, 66%). This was especially pronounced in the grasses, where we found evidence for phylogenetic signal for cell aspect ratio in 7 out of 8 (88%) sampled grasses (Fig. 3c). The gymnosperms also exhibited phylogenetic signal for aspect ratio (Fig. 3d). In the eudicots, evidence for phylogenetic signal in both traits was concentrated in closely related species. There was no strong evidence for particular shape metrics characterizing families, orders, or other major eudicot clades. Thus, while eudicot epidermal cell shapes were not distinguished by particular shape metrics, fern epidermal cells are characterized by high undulation, core monocot epidermal cells by low undulation and low aspect ratio, and gymnosperm cells by low aspect ratio.

**Figure 3.**
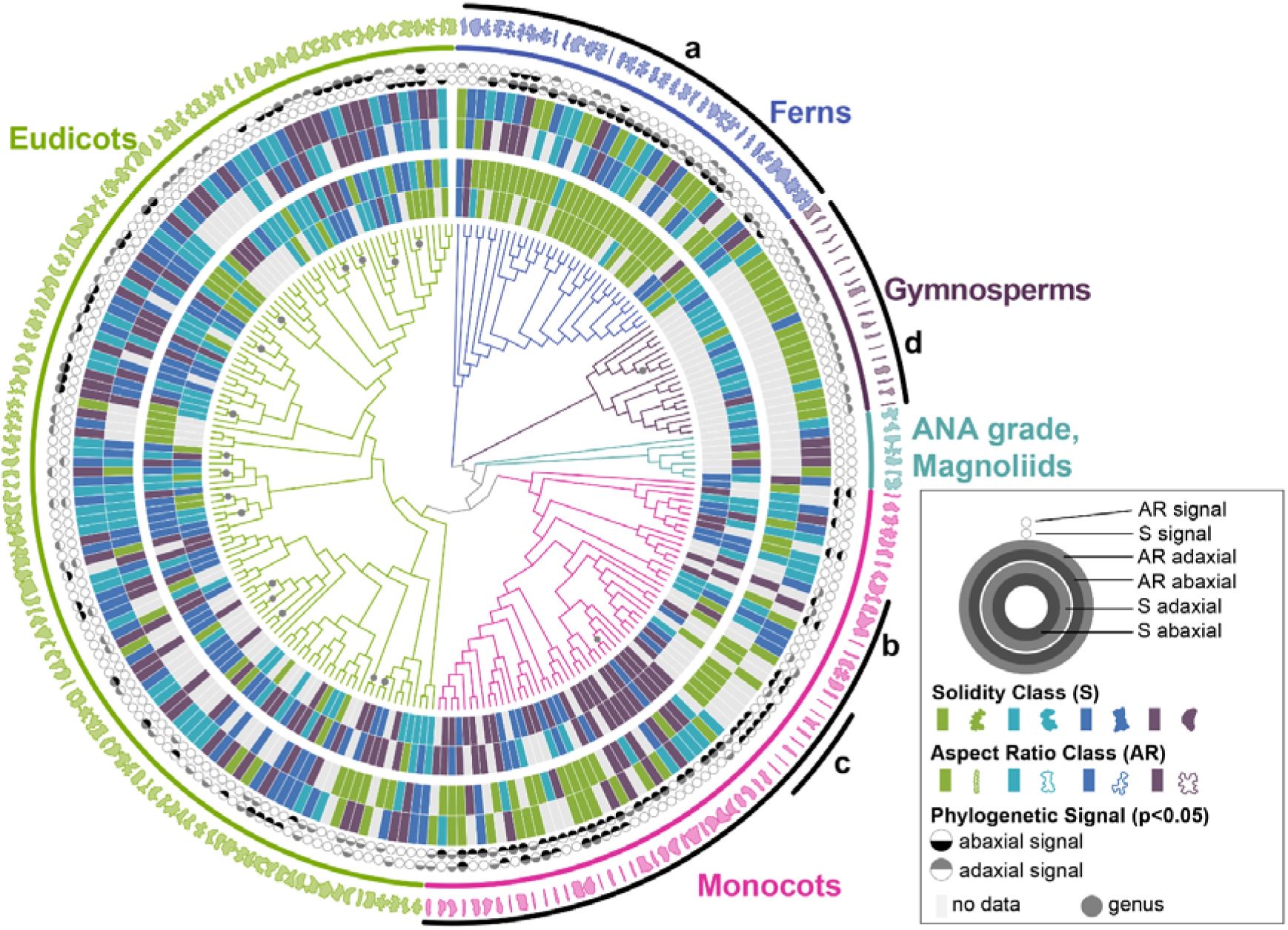
Pavement cells in the ferns, gymnosperms, and monocots are characterized by particular shape metrics, while eudicots exhibit wide variation. A phylogenetic reconstruction of taxa sampled in our dataset (centre) surrounded by data rings depicting cell aspect ratio and solidity, and leaf aspect ratio (see legend for positional key). Branch lengths are not shown in this figure, although they were used in all analyses. Taxonomic groups are indicated by colour. One representative cell shape from each species is depicted on the outermost ring. Specific clades are indicated as follows: (a) ferns; (b) the core monocots; (c) the grasses; (d) gymnosperms. Each grey dot on tree indicates multiple species in the same genus. All data and species names can be found in Table S1.

This result suggests that in the ferns and in the core monocots, aspects of either the cell margin patterning machinery (e.g. actin and microtubule dynamics), or wall material properties, are shared between members of each clade. In the eudicots, these cell shape generating mechanisms and cell wall properties may be more variable at large evolutionary distances. However, trait values between closely related species (e.g. species in the same genus) were often correlated, even in the eudicots (Fig. 3). Indeed, epidermal cell traits can be used as characters in systematics studies^47–49^. This highlights the critical importance of accounting for phylogeny when testing hypotheses about the function of particular epidermal cell shapes^20^. For example, particular epidermal cell shapes have been proposed to be important in drought tolerance, in focusing light onto the photosynthetic machinery, or in providing mechanical stability to the epidermis^50–52^. When these hypotheses are tested using multiple different species, it is important to remember that cell shapes may be similar between species not because of a particular function, but because of underlying phylogeny.

### Abaxial leaf surfaces present more undulate cells

Sparse qualitative observations indicated that abaxial cells tended to have more undulate margins than adaxial cells^43^. To test this across our sampling, we calculated the difference between the average adaxial solidity and the average abaxial solidity in the 146 species for which we had data from both sides of the leaf (81 eudicots, 30 monocots, 28 ferns; see Methods). We found that when a difference in cell solidity was present, the abaxial cells tended to have more undulations (lower solidity, Fig. 2a), in line with the qualitative literature^16,53^. The causes of such differences are ripe for discovery. In many cases, different sides of the leaves experience different microclimates; undulation exhibits some environmental plasticity and thus it is plausible that more undulation on abaxial surfaces could relate to local environmental influences^43,54–56^. In addition, abaxial vs. adaxial developmental identity may contribute to differential undulation^43,57,58^. The number of cells of other cell types on the abaxial surface, particularly increased stomatal number^59^, could contribute to increased undulation through a packing adjustment. Lastly, it is possible that differential growth rates between the two sides may relate to differential undulation; however, such growth differences would need to be balanced to finally yield a flat leaf. In *curl* tomato mutants, whose curled leaves exhibit larger cells on the abaxial epidermis, there are no qualitative differences in abaxial (or adaxial) cell undulations from the wild type^60^.

### Anisotropic leaves tend to have anisotropic cells

We next wanted to explore connections between leaf shape and cell shape. Final epidermal cell shape over the surface of a leaf is a record of the developmental history of growth patterns – highly anisotropic cells indicate directional cell expansion, while regions of smaller cells indicate cell expansion coupled with division^61,62^. Leaf form is likely generated by complex growth patterns that we would be unable to detect with our sampling^63^. However, in the Brassicaceae, a connection between growth direction, cell shape, and organ shape has recently been proposed^19^. In addition, in the flowers of *Saltugilia spp*.^64,65^ and *Mimulus lewisii^66^*, highly anisotropic epidermal cells are present on anisotropic floral tubes, and more isotropic cells on petal lobes. We wondered whether we would be able to detect a similar connection between cell aspect ratio and leaf aspect ratio at the broad scale of our dataset.

To explore any connection between leaf and cell aspect ratio in our sampling, we examined correlations between leaf aspect ratio and cell aspect ratio. We found evidence for a correlation between leaf and cell aspect ratio: highly anisotropic leaves tended to have highly anisotropic cells, but not in the eudicots (Fig. 4a). This indicates that in the anisotropic leaves of some vascular plants, anisotropic growth likely involves cell expansion, with very little coincident cell division, across large regions of the growing leaf. Were the cells to divide as they expanded, this connection would have been lost. In contrast, even in eudicots with leaves in the lowest aspect ratio quartile, cell aspect ratio levels never reached the lowest quartile. For example, in *Plantago afra* (eudicot) and *Hemerocallis fulva* (monocot), leaf aspect ratio values were similar (0.091 and 0.087, respectively. Both class 4), but mean cell aspect ratio values were not (mean AR_pa_ = 0.67, class 3; mean AR_Hf_ 0.17, class 1). In the sampled eudicots as a whole, we found no correlation between cell aspect ratio and leaf aspect ratio. This indicates that growth dynamics differ considerably between different clades, even when leaf form is superficially similar^67^. While cell division and elongation are essential drivers of growth and development, the development of plant form can only be understood by studying the balance between these two processes, and their regulation in an organ-level and organismal context^62,68–70^.

**Figure 4.**
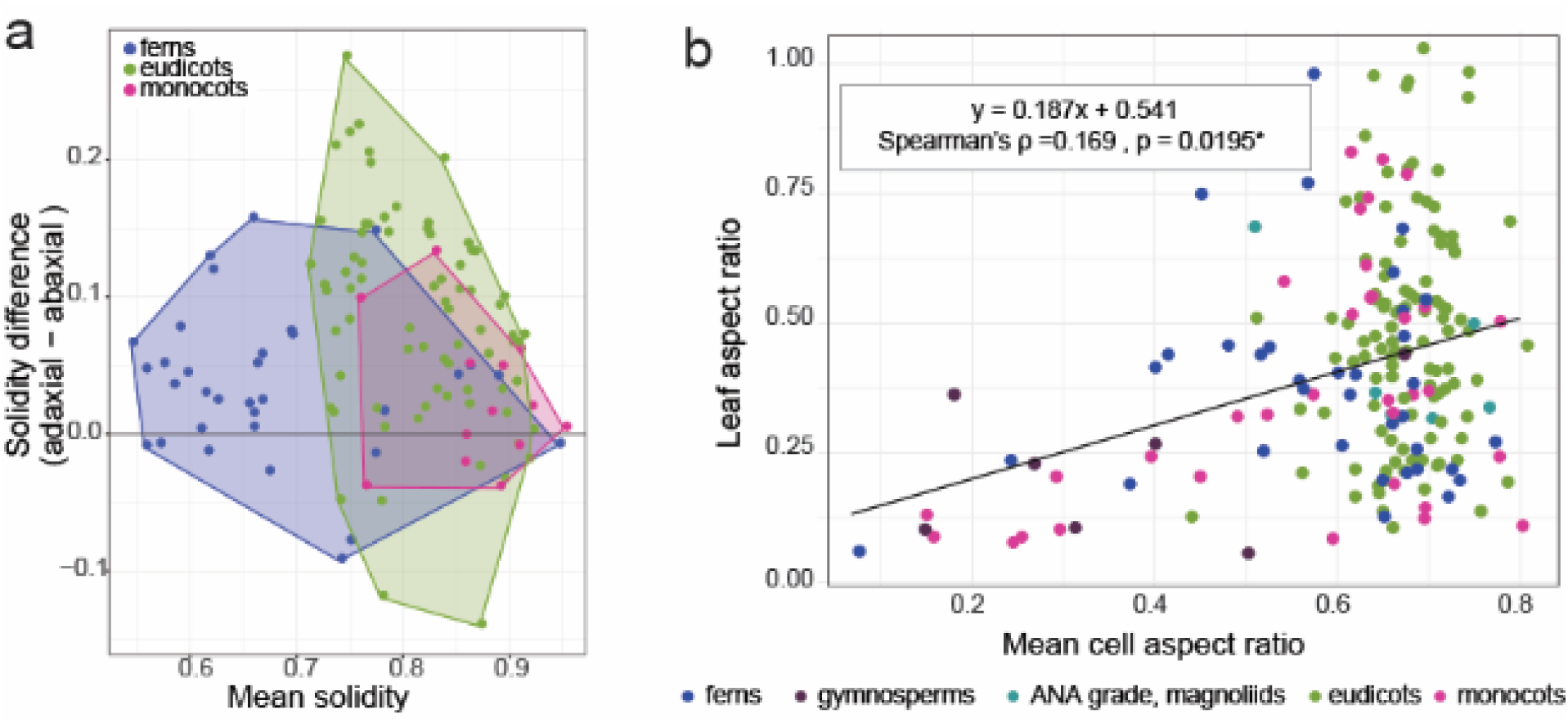
Cell metrics vary according to particular leaf traits. (a) Cell solidity is often lower (higher undulation) on the abaxial leaf face. (b) In the full data set, regardless of leaf side, mean cell aspect ratio and mean leaf aspect ratio are correlated. Anisotropic leaves tend to have anisotropic cells, but not in the eudicots. Linear regression line, based on all data, is shown with Spearman’s (ρ) correlation coefficient. All data can be found in Table S1.

A second connection between cell shape and organ form that has been proposed is that the highly undulating cells characteristic of some eudicots are a consequence of cell expansion in all directions in the plane of the leaf lamina^19,58,71^. In this case, one would expect highly anisotropic leaves to have cells with high solidity values; or that highly anisotropic cells would have high solidity values (fewer undulations). We detected no correlations between cell solidity and leaf aspect ratio, or between cell solidity and cell aspect ratio (Fig. S2). Thus, while margin undulation may not be related to organ shape in our broad sample, in non-eudicot species there is a correlation between low leaf aspect ratio and low cell aspect ratio. Further taxonomically broad exploration of cell expansion and division over time, similar to that applied in arabidopsis^61,72,73^, would prove highly informative for understanding the breadth of organ growth mechanisms present in the plant kingdom.

### Conclusions

Our analysis has revealed striking diversity in leaf epidermal cell shape. This quantitative analysis has allowed for mapping of shape metrics in a phylogenetic context, demonstrating that while closely related eudicots tend to share cell shape characters, there is no obvious global trend of trait retention in this clade. The lack of consistent highly undulatory cell margins, like those observed in arabidopsis, make a strong case for expansion beyond a single model system. Similarly, while maize epidermal cells have highly undulate margins, monocots show a phylogenetic signal for weakly undulating cells, again pointing to a need to work in species beyond the grasses.

How might epidermal shape diversity arise? Based on the well-resolved molecular network regulating cell shape in arabidopis (and maize)^2–4,6,9,74^, an attractive hypothesis might be that the patterning system, centred on ROP-mediated exclusivity between actin and microtubule position, is variable among species. Variability in the patterning of cell wall synthesis and modification would yield variation in cell undulation. Alternatively, the cytoskeletal patterning mechanism might be perfectly consistent in most species (suggested by conservation between arabidopsis and maize), but cell wall synthesis and modification might differ between species and clades. Indeed, primary cell wall composition is highly variable across plants^75^. Looking to diversity in a quantitative phylogenetic framework will be critical in determining both how diverse cell shapes arise, and what their functions might be within organs and organisms.

## Acknowledgements

The authors thank Joanna Wolstenholme, Jeffrey Heithmar, and Rebecca Goldberg, for preliminary work and technical assistance. We also thank the community attending FASEB Plant Development, a meeting at which this collaboration began. Tobias Baskin provided helpful comments on an earlier version of the manuscript. The University of Cambridge Botanic Garden and the UMass Natural History Collections are thanked for access to their collections, with special thanks to Sally Petit (CUBG), Christopher Phillips (UMass), Daniel Jones (UMass), and Alex Summers (CUBG). Work in the Braybrook group is supported by UCLA and previously was supported by The Gatsby Charitable Foundation (GAT3396/PR4) and R.V was supported by the EPSRC Cambridge NanoDTC (EP/G037221/1). Work in the Bartlett group is supported by the UMass Biology Department (G.D.P) and the NSF CAREER program (IOS-1652380).

## Author Contributions

S.B. conceived of the original project and designed experiments with R.V. and M.B. M.B. and G.D.P. devised a sampling strategy. R.V and G.D.P collected samples, prepared and imaged samples, generated outlines, and utilised momocs for analyses. R.V, J.G., and M.B. performed in depth analyses with momocs, generated PCAs, and examined correlations between descriptors. M.B. performed phylogenetic signal analyses. S.B. and M.B wrote the manuscript and prepared figures, with assistance from R.V. and J.G.

---

**Figure S1.**
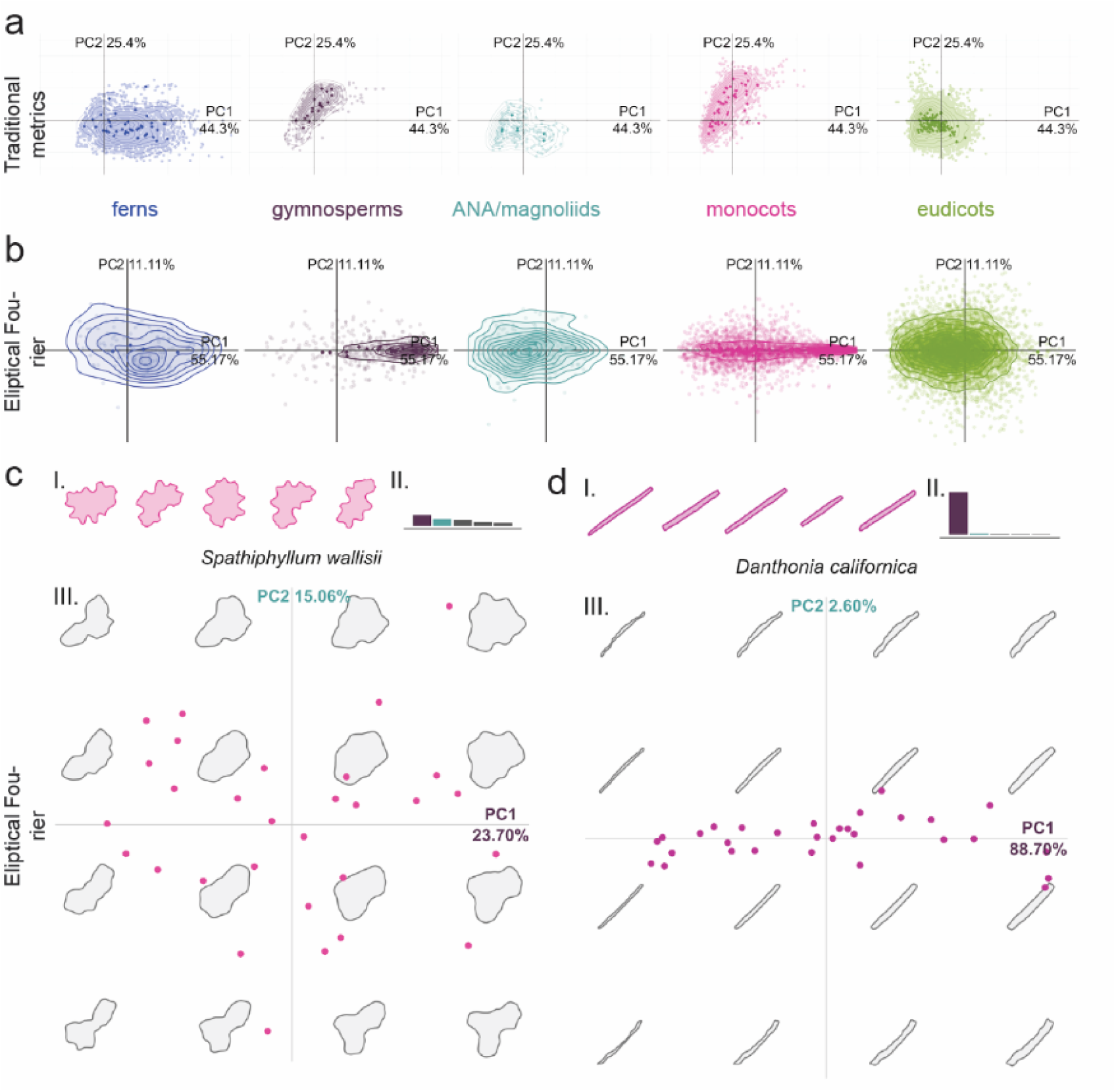
Results from Traditional and Elliptical Fourier analysis of cell shapes. Morphospace for PCA1 and PCA2 (a) Traditional; (b) Elliptical Fourier, split by clade. The analysis was performed on all groups together but shown here independently. Fourier analysis morphospaces for two monocot examples (c) *Danthonia californica* and (d) *Spathiphylium wallisii*, exhibiting margin undulation and cell anisotropy, respectively. Representative cells shapes (I), eigenvalues for PCA1 (purple) and PCA2 (turquoise) (II), and PCA1 and PCA2 morphospaces (III); this analyses demonstrates the utility of Elliptical Fourier analysis for cell base shape (anisotropy in *Danthonia* is well described by 88.7% in PCA1) and its lacking when margin undulation is the variable trait (*Spathiphyllum*, low percentage of variation explained by any one PCA as show in eigen values).

**Figure S2.**
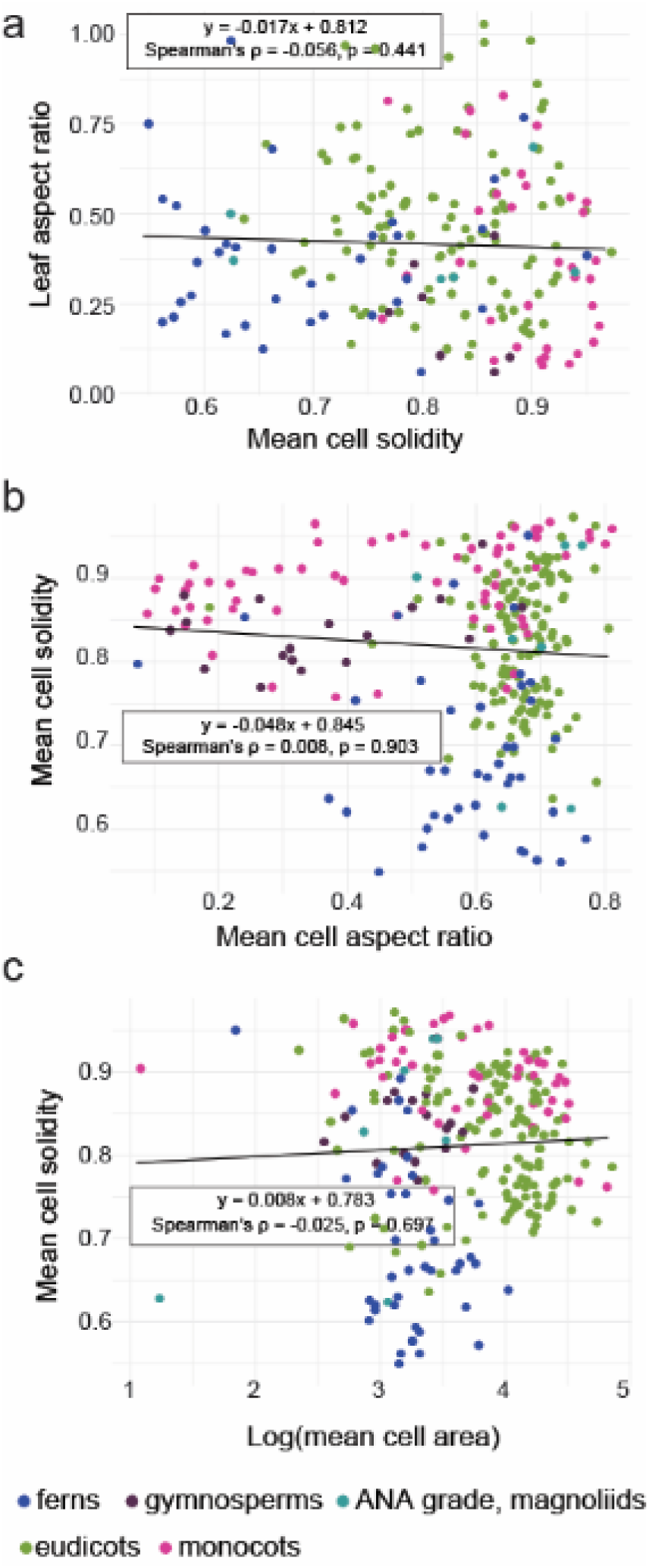
Correlation plots for cell and leaf metrics. No correlation was evident between (a) mean cell solidity and leaf aspect ratio; (b) mean cell aspect ratio and mean cell solidity; (c) log(mean cell area) and mean cell solidity.

**Figure S3.**
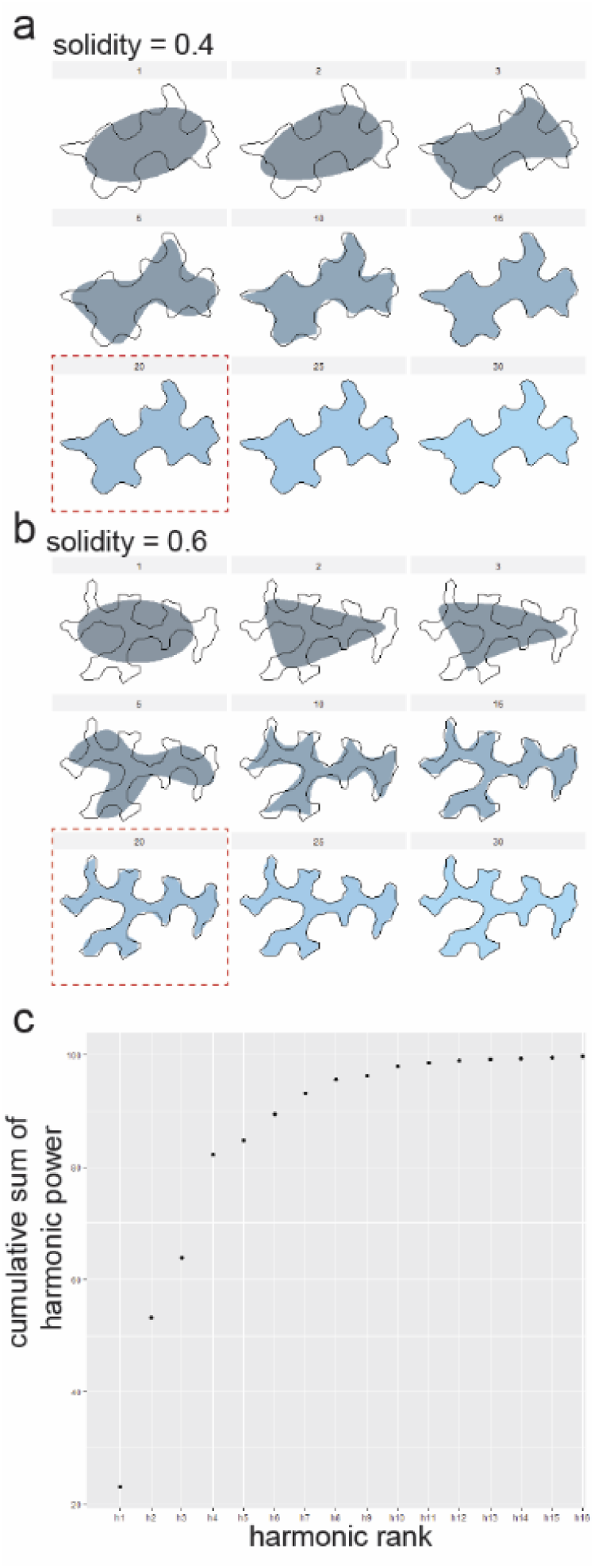
Harmonic assessment for Elliptical Fourier Analysis. Representative cell shapes of solidity values 0.6 (a) and 0.4 (b) with fitted ellipses at several harmonic ranks. While the less undulatory cell (S=0.6) could be fit with 15 harmonics, the more undulatory (S=0.4) cell is best fit by 20 harmonics. For this reason, 20 harmonics were chosen for our analyses. (c) a graph of the cumulative sum harmonic power for our dataset (a measure of how many harmonics are required to fit the data) and 99.9% is easily reached with h16.

